# MyoL-Dependent Coupling of Preconoidal Rings to Conoid Is Required for Motility in *Toxoplasma gondii*

**DOI:** 10.64898/2026.01.06.697858

**Authors:** Romuald Haase, Albert Tell i Puig, Nicolas Dos Santos Pacheco, Bohumil Maco, Oscar Vadas, Dominique Soldati-Favre

## Abstract

Apicomplexan parasites are unified by their apical complex, a structure of cytoskeletal elements and secretory organelles. At its core lies the conoid, made of spiraling tubulin fibers essential for parasite motility and invasion. In *Toxoplasma gondii*, conoid extrusion and retraction through the apical polar ring is powered by Myosin H, ultimately regulating gliding motility. The conoid is capped by three preconoidal rings (PCRs), critical for extrusion and motility as they anchor formin 1, which produces filamentous actin needed for both processes.

We demonstrate here that PCR composition and stability differ between daughter and mature cells and that Pcr2 and Pcr3 adopt a half-ring localization. The non-functional head containing Myosin-like protein MyoL, a previously unrecognized PCR component, is essential for gliding motility and invasion. Its depletion partially detaches PCRs from the conoid, highlighting its role as a mechanical tether. Targeted mutagenesis identifies the neck and tail regions of MyoL as critical for localization and function. Together, these findings illuminate the architecture and functional organization of the PCR in *T. gondii*, a structure conserved across several Apicomplexa.

## Introduction

The Apicomplexa phylum groups species of high veterinary and human health importance such as *Plasmodium spp*, *Cryptosporidium spp* and *Toxoplasma gondii,* respectively causative agents of malaria, cryptosporidiosis and toxoplasmosis (Adl et al., 2007; Guérin & Striepen, 2020; Montoya & Liesenfeld, 2004; Phillips et al., 2017). Members of this phylum are characterized by the presence of an apical complex, composed of cytoskeletal elements and secretory organelles (Dos Santos Pacheco et al., 2020). Within the apical complex stands the conoid, a cytoskeletal structure made of spiraling tubulin fibers and topped by three proteinaceous preconoidal rings (PCRs) (Gui et al., 2023; Hu et al., 2002). Additionally, two short microtubules, termed intraconoidal microtubules (ICMTs), reside inside the cone **(Fig 1A)**. The conoid is a dynamic structure that moves through the apical polar ring (APR) during motility and invasion in response to intracellular calcium waves (Del Carmen et al., 2009; Mondragon & Frixione, 1996; Monteiro et al., 2001). The extrusion and retraction of the conoid have been shown to regulate parasite motility by directing F-actin generated at the PCRs into the pellicular space, making it accessible to the glideosome, the motor machinery that propels the parasite forward (Dos Santos Pacheco et al., 2022; Frénal et al., 2010; Martinez et al., 2023; Ren et al., 2024). Through this process, the conoid functions as a structure finely tuning parasite motility and invasion (R. Haase et al., 2022).

**Figure 1.**
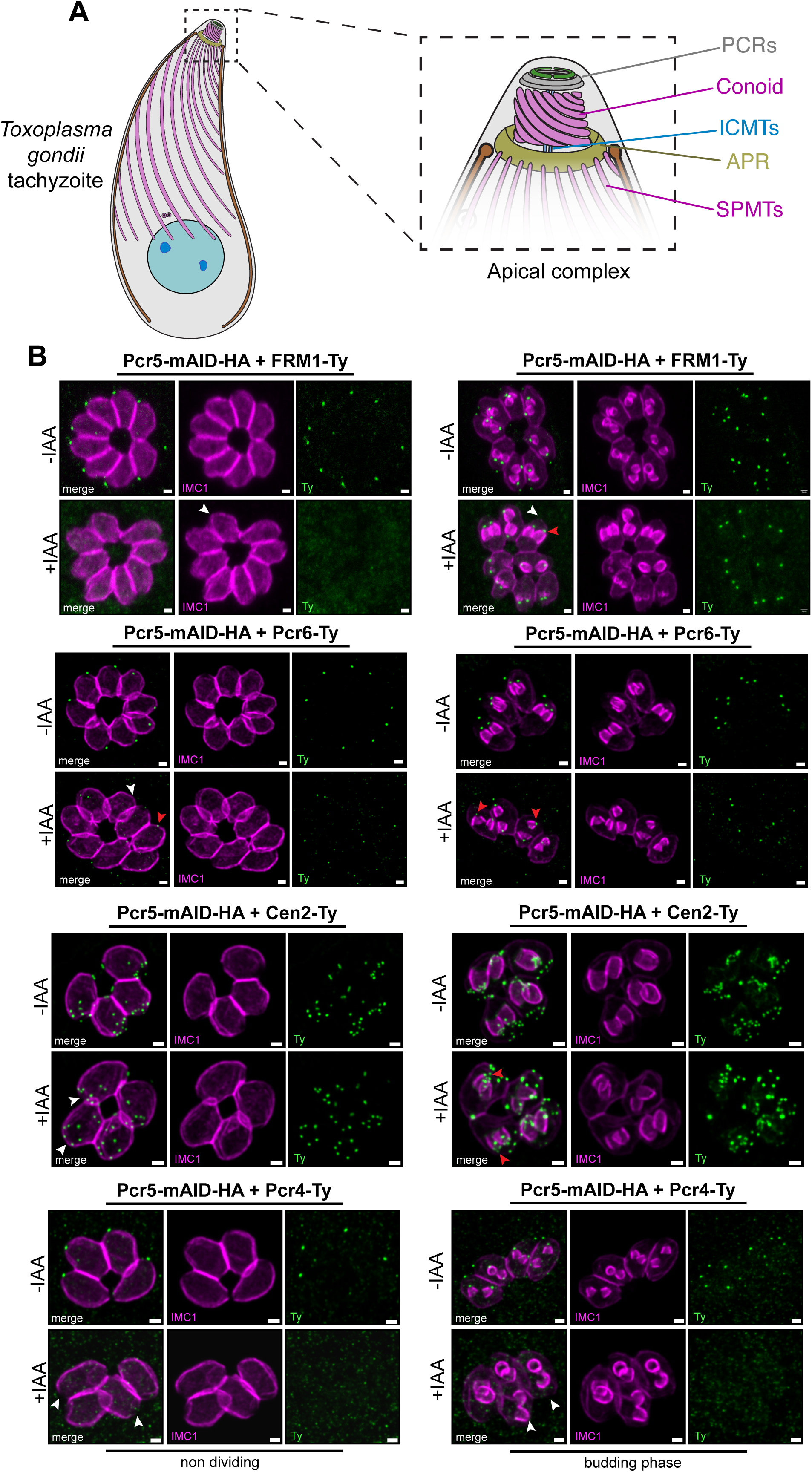
Components of the PCRs exhibit differential stability between daughters and mother cells. **(A)** Schematic representation of the cellular architecture of *T. gondii* tachyzoites. Right panel, zoom in the apical complex components. **(B)** IFA images of FRM1, Pcr6, and CEN2 in mature intracellular parasites and during daughter cell formation in the presence or absence of Pcr5. Scale bar = 1µm. Red arrowheads indicate when the signal is present, whereas white arrowhead indicates its absence.

The PCRs play a central role in this mechanism by anchoring key glideosome components, including formin 1 (FRM1), which nucleates F-actin at the apical end, and the Glideosome-Associated Connector (GAC), which links the translocating actin filaments to the parasite plasma membrane (Daher et al., 2010; Dos Santos Pacheco et al., 2022; Jacot et al., 2016; Tosetti et al., 2019). The PCRs are notably composed of proteins essential for their integrity as parasites lacking Pcr4 or Pcr5 lose their PCRs, which in turn impairs their ability to disseminate and invade host cells (Dos Santos Pacheco et al., 2022). In addition, Pcr6 functions as a molecular anchor, tethering the PCRs to the cone of tubulin fibers (Dos Santos Pacheco et al., 2022). The function of some PCR components remains elusive, such as Pcr2, whose depletion severely impairs egress and invasion without affecting the overall PCRs structure (Munera Lopez et al., 2022). Despite these insights, little is known about the biogenesis of the PCRs and the other apical complex elements. Detailed microscopy studies have shown that the first element to be formed is the APR, followed by the conoid fibers nucleated by γ-tubulin, and finally the PCRs (Arias Padilla et al., 2024; Engelberg et al., 2025; Haase et al., 2024; Padilla et al., 2024). Nonetheless, a molecular characterization of the proteins taking part in the formation of the PCRs and its contribution remains to be elucidated. Here, we conducted a detailed investigation into the composition of PCRs in daughter cells. We found that parasites depleted for Pcr5 and lacking PCRs in mature cells still maintain certain PCR markers at the tips of daughter cells, indicating differential stability of the structure among the different developmental stages. Furthermore, Ultrastructure Expansion Microscopy (U-ExM) revealed that Pcr2 and Pcr3 form an atypical “half-ring” staining pattern, similar to what was previously described for Pcr1 (Dos Santos Pacheco et al., 2022; Koreny et al., 2021). Importantly, we identified myosin-like protein MyoL as a novel PCRs component. Parasites depleted of MyoL display severe defects in gliding motility and host cell invasion, attributable to the partial detachment of PCRs from the conoid. Targeted mutagenesis demonstrated that the neck-tail region of MyoL is essential for its correct localization and function while the non-functional motor domain is dispensable. Together, these findings offer new insights into the composition and stability of the PCRs as well as its assembly to the conoid, a structure conserved across Apicomplexa and critical for parasite pathogenicity.

## Results

### Upon depletion of Pcr5, the preconoidal rings are only lost in mature parasites

We previously identified two components of the PCRs, Pcr4 and Pcr5, which appear early during daughter cell formation and are essential for the stability of the PCRs in mature parasites (Dos Santos Pacheco et al., 2022) **(Fig 1A)**. However, it was unclear whether the loss of PCRs following Pcr4 or Pcr5 depletion resulted from impaired biogenesis or from disassembly in mature cells. The PCRs are among the earliest structures formed during endodyogeny (Padilla et al., 2024) To investigate their biogenesis, we monitored by indirect immunofluorescence assay (IFA), the PCRs markers FRM1, Pcr6, Cen2 and Pcr4 (Dos Santos Pacheco et al., 2022) in the daughter cells **(Fig 1B)**. As previously observed, Pcr5 depletion leads to the disappearance of apical FRM1, Pcr4 and Cen2 proteins in mature intracellular parasites while some parasites maintained a Pcr6 staining at their apical tip (Dos Santos Pacheco et al., 2022) **(Fig 1B)**. However, during the budding phase, when the IMC attaches to the growing daughter cell cytoskeleton, all the daughter cells displayed apical staining for FRM1, Pcr6 and Cen2 in the Pcr5 depleted parasites **(Fig 1B)**. In contrast, Pcr4 known to interact as an heterodimer with Pcr5, could not be detected at the tip of daughter cells (Dos Santos Pacheco et al., 2022) **(Fig 1B)**.

These observations suggest that, in the absence of Pcr5, the PCRs are built and correctly positioned during daughter cell formation but are lost at a late stage of endodyogeny. Pcr4 that forms a heterodimer with Pcr5 remains the only dramatically impacted protein at early stages. This suggests that the Pcr4/Pcr5 heterodimer is dispensable for PCRs formation but essential for their stability.

### Pcr2 and Pcr3 form half-rings and display atypical behavior upon loss of the PCRs

To further investigate the stability of the PCRs, we monitored two previously described PCRs components, Pcr2 and Pcr3 (Munera Lopez et al., 2022). These proteins were endogenously Ty-tagged at their C-terminal side in the Pcr5-mAID-HA strain and correct epitope-tagging was verified by western-blot **(Fig S1A)**. Unexpectedly, despite the consistent loss of apical FRM1 staining in Pcr5-depleted mature parasites, reflecting PCRs disassembly, Pcr2 and Pcr3 remained apically localized in mature parasites **(Fig 2A)**. IFA on extracellular parasites showed the same phenotype, with Pcr2 and Pcr3 remaining at the apical pole **(Fig 2B)**. To determine their association with the PCRs, Pcr2 and Pcr3 were endogenously tagged with a mAID-HA cassette and correct integration was verified by genomic PCR **(Fig S1B)**. U-ExM revealed that both Pcr2 and Pcr3 exhibit a characteristic "half-ring" staining pattern **(Fig 2C)**. To investigate further their positioning, we endogenously epitope-tagged the Pcr1, previously reported as a “half-ring” (Dos Santos Pacheco et al., 2022), into the Pcr2-mAID-HA and Pcr3-mAID-HA strains, which were verified by western blot **(Fig S1C)**. Colocalization of Pcr1 with Pcr2 and Pcr3 showed perfect overlap in their vertical positioning **(Fig 2D)**. However, while Pcr1 and Pcr2 localized to the same side of the structure, Pcr3 was observed on the opposite side of Pcr1, effectively reconstituting a full ring when the two signals were overlaid **(Fig 2D)**.

**Figure 2.**
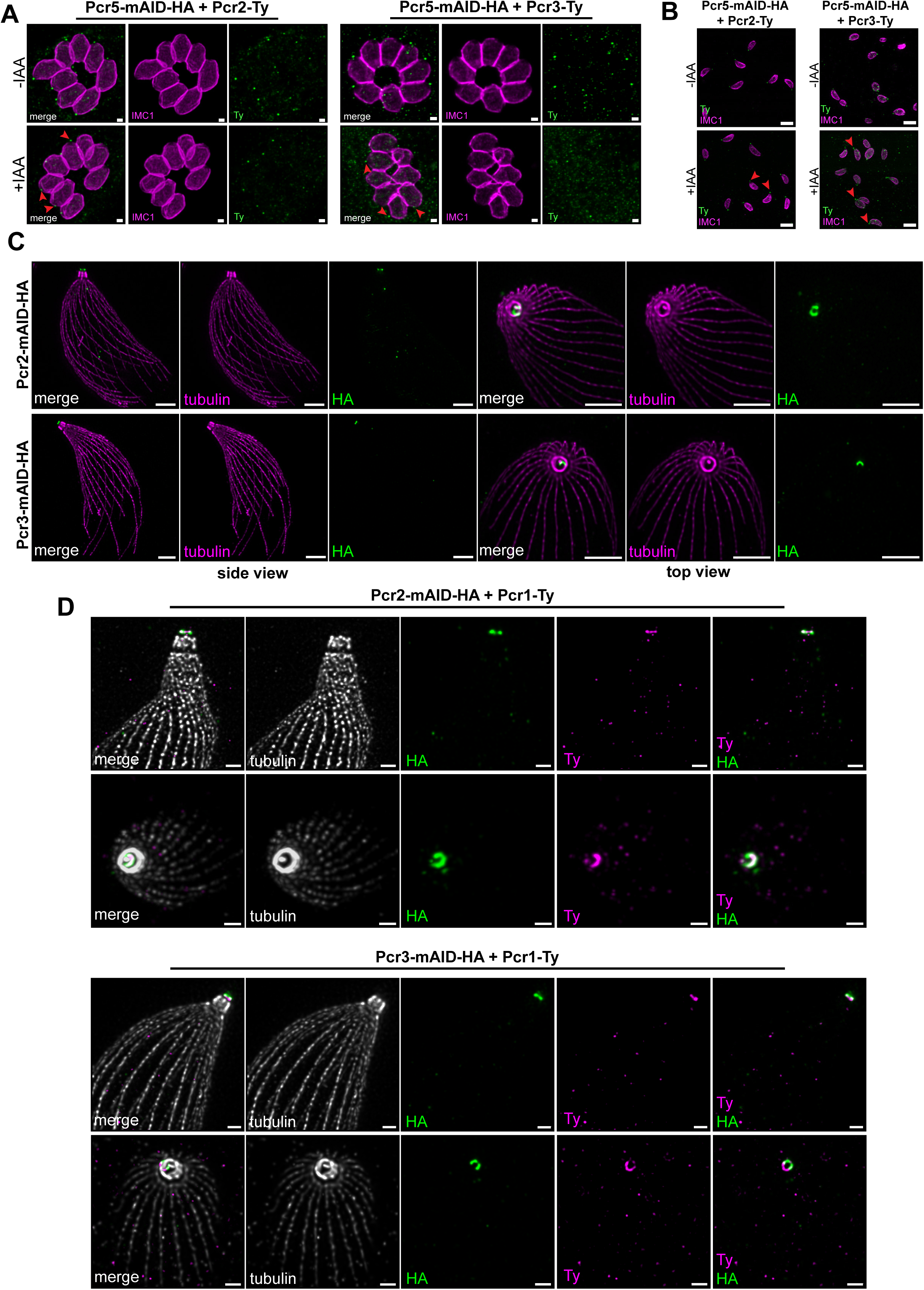
Pcr2 and Pcr3 are half-ring PCRs and behave independently of Pcr5. **(A)** IFA images of Pcr2 and Pcr3 in mature intracellular parasites in the presence or absence of Pcr5. Scale bar = 1µm. **(B)** IFA images of Pcr2 and Pcr3 in extracellular parasites in presence or absence of Pcr5. Scale bar = 5µm. **(C)** U-ExM localization of Pcr2 and Pcr3 proteins in extracellular parasites in side and top view. Scale bar = 3µm. **(D)** Colocalization in U-ExM of Pcr1 with the Pcr2 and Pcr3 proteins in side and top view. Scale bar = 3µm.

A putative third PCR structure was recently identified by cryo-electron tomography (cryo-ET (Gui et al., 2023), which may correspond to a plasma membrane-associated rather than a conoid-linked element. To test whether Pcr1, Pcr2, and Pcr3 remain associated with the detached conoid complex or instead stay with the apical cell body, we used the RNG2-mAID-HA strain. In parasites depleted of RNG2, the conoid complex, including the PCRs and the ICMTs, detach from the apical pole upon parasite activation, during the retraction of extruded conoid (Haase et al., 2025) **(Fig 3A)**. Pcr1-2-3 were endogenously epitope-tagged into the RNG2-mAID-HA strain and successful tagging was verified by western blot **(Fig S1D)**. All three proteins localized above the tubulin cone in the absence of auxin. Upon auxin treatment, they remained associated with the detaching conoids, as evidenced by neighboring staining **(Fig 3B)**. Taken together, these data suggest that Pcr1-2-3 are intrinsic components of the PCRs that form a stable structure that remains apical upon either PCRs dissociation or conoid detachment.

**Figure 3.**
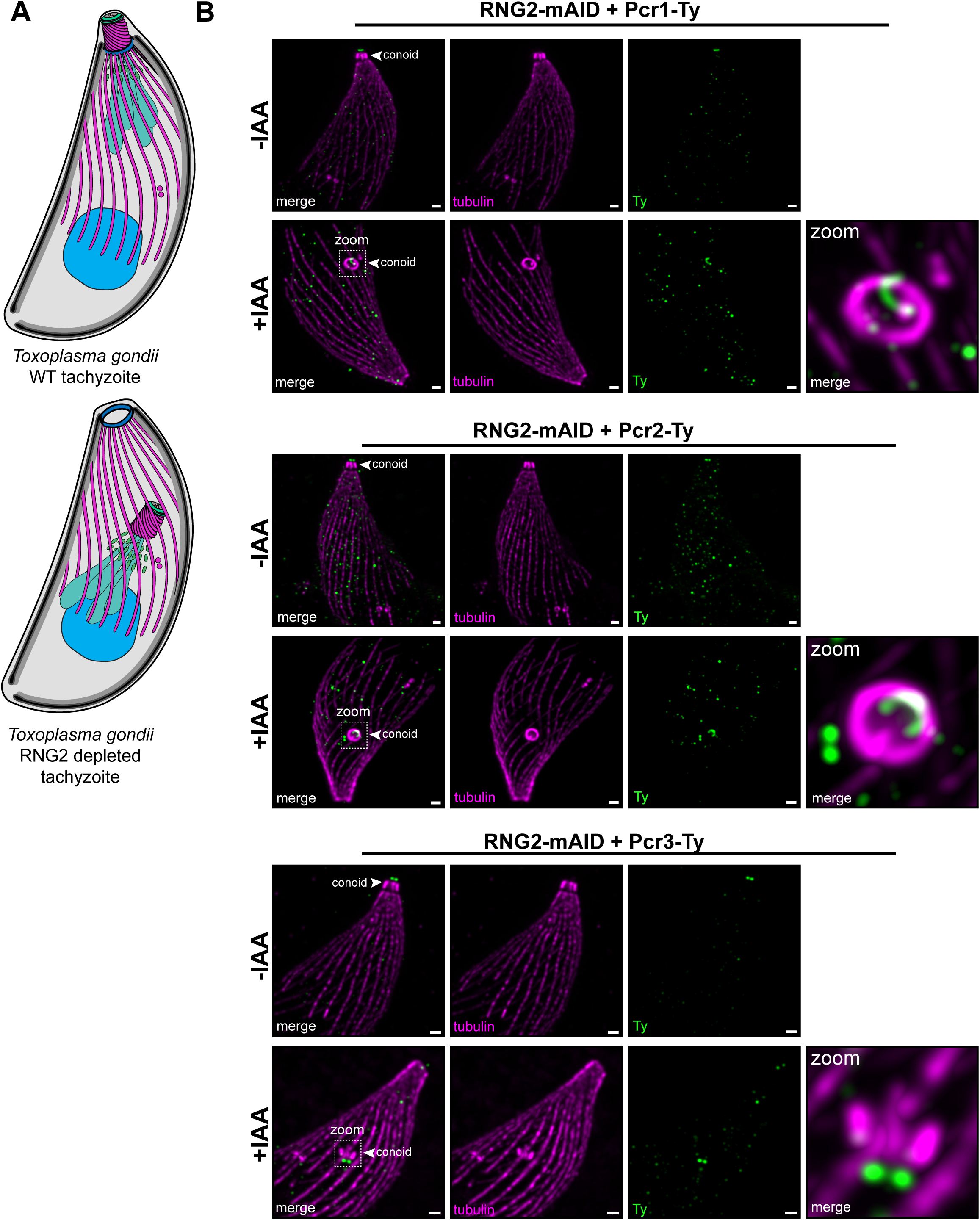
Pcr2 and Pcr3 remain associated with the detached conoid in RNG2 depleted parasites. **(A)** Schematic representation of the phenotype upon RNG2 depletion in *T. gondii* tachyzoites. **(B)** U-ExM localization of Pcr1, Pcr2 and Pcr3 proteins in extracellular parasites depleted or not for the RNG2 protein. Scale bar = 1µm.

### MyoL is a non-functional myosin-like protein with a divergent RCC1-like domain

Among the proteins localized to the apical tip but not yet assigned to a specific structure by U-Ex-M is MyoL. It localizes both in the cytoplasm and as a small apical focus (Frénal et al., 2017) and it is predicted to be fitness conferring (Sidik et al., 2016). MyoL is a myosin-like protein that structurally resembles a myosin but lacks the key features of an active motor. Sequence analysis reveals an N-terminal loop region followed by a head-like domain, an α-helical neck containing four IQ motifs, and a C-terminal RCC1 domain suggesting structural instead of motor function **(Fig 4A)** (Foth et al., 2006; Heintzelman & Schwartzman, 1997). Although MyoL is predicted to adopt a head domain conformation similar to MyoH or human myosin-V **(Fig 4B)**, its ATP binding site and actin binding region are highly divergent **(Fig 4C-D)** (Pospich et al., 2021). Additionally, the head domains of MyoH and MyoA align very well, whereas the head domain of MyoL shows poor alignment, further supporting its divergence from other myosins **(Fig S2A-B)**. This suggests that MyoL is a non-functional motor, which however retains a neck domain with IQ motifs capable to associate with Myosin Light Chains (MLCs). Recently published Cryo-EM structure of the preconoidal ring 2 identified three light chains associated with MyoL (MLC4, Calmodulins 1 & 2), supporting the findings that MyoL and its associated light chains have a structural function at the PCRs (pdb: 9YA0) (Jianwei Zeng, 2025). Similar to MyoH and other myosins, its stability and correct folding require UNC, a myosin-specific co-chaperone of the UCS (UNC-45/CRO1/She4p) family (Frénal et al., 2017). The MyoL RCC1 domain, which is absent from the PCR-P2 ring Cryo-EM structure, contains several insertions between the blades of the β-propeller compared to MyoH, which may enable unique binding interactions (pdb: 9YA0) (Jianwei Zeng, 2025) **(Fig 4E)**. Blast-p analysis revealed orthologs of MyoL in several apicomplexan parasites including *Plasmodium* and *Cryptosporidium species* **(Table 1)**. To reflect its divergence from other myosins while maintaining consistency with the established nomenclature we refer here to Myosin-Like (MyoL). Overall, MyoL is a conserved myosin-like protein which contains characteristic features reminiscent of conventional myosins with unique insertions, making this protein a non-functional divergent myosin.

**Figure 4.**
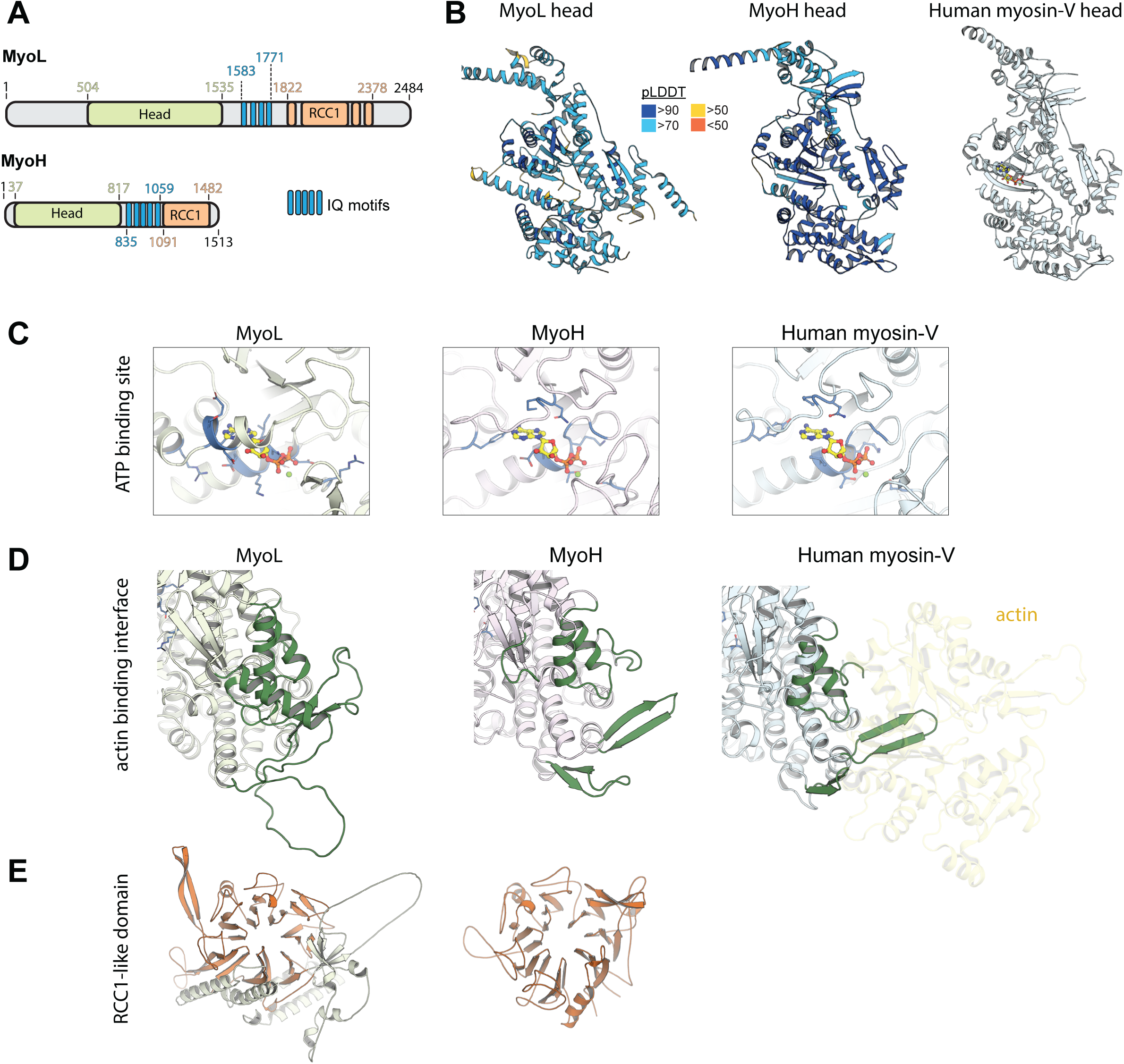
MyoL is structurally related to MyoH and harbours a non-functional motor domain. **(A)** Domain organization of MyoL and MyoH. Boundaries were identified combining sequence and structural information **(B)** Comparison of predicted head domain conformations of MyoL and MyoH with the experimentally determined structure of human myosin-V (pdb: 7pm9). Only high-confidence regions (pLDDT score > 50) of the predictions are shown and colored according to legend. **(C)** Zoomed-in views of the ATP-binding sites. ADP position was modelled for MyoL and MyoH based on alignment with myosin-V complex structure. **(D)** Zoomed-in views of actin-binding region, with residues directly mediating actin binding colored in green. **(E)** Homo-dimerisation prediction of the neck-tail constructs for MyoL and MyoH colored as in panel A. **(F)** Comparison of the RCC1 domain structure prediction of MyoL and MyoH highlighting the large insertions in MyoL.

### MyoL is PCRs component contributing to conoid tethering

To investigate its localization and function, MyoL was endogenously tagged with a mAID-HA cassette at its C-terminal site (Brown et al., 2018; Dos Santos Pacheco & Soldati-Favre, 2021). Correct integration and clonality were assessed by PCR on genomic DNA **(Fig S3A)**. As previously reported, MyoL localizes as an apical dot at the parasite tip and U-ExM further revealed a ring-like pattern above the conoid, supporting its association with the PCRs **(Fig 5A – S3B)**, throughout all the stages of endodyogeny **(Fig S3C)**. Western blot confirmed efficient degradation upon 24h of auxin (IAA) treatment **(Fig 5B)** and depletion of MyoL for seven days resulted in a marked reduction in plaque size, indicating a significant defect in the parasite’s lytic cycle **(Fig 5C-D)**. While MyoL depletion severely compromised host cell invasion **(Fig 5E)**, parasites only showed a minimal reduction in their capacity to egress, successfully lysing the parasitophorous vacuole membrane (PVM) in the induced egress assay **(Fig 5F)**. Nevertheless, despite successful PVM lysis, MyoL-depleted parasites exhibited a clear impairment in spreading to adjacent host cells, as revealed by the egress assay **(Fig 5G)**. To validate these findings, a gliding trail assay was conducted. While untreated parasites generated abundant linear trails, consistent with efficient motility and dissemination, MyoL-depleted parasites displayed aberrant circular trails, demonstrating a defect in their gliding-mediated spread **(Fig 5H)**. To further validate the gliding motility defect, we performed live imaging microscopy of induced egress parasites in presence or absence of MyoL, showing a clear defect in motility upon MyoL depletion **(Supplementary Movie 1-4)**. Moreover, parasites depleted from MyoL had a 50% reduction in conoid extrusion **(Fig 5I)**. The observed gliding motility defect was not due to a microneme secretion defect, since MyoL-depleted parasites exhibited normal microneme release, in agreement with their ability to effectively undergo induced egress **(Fig 5J – S3D)**.

**Figure 5.**
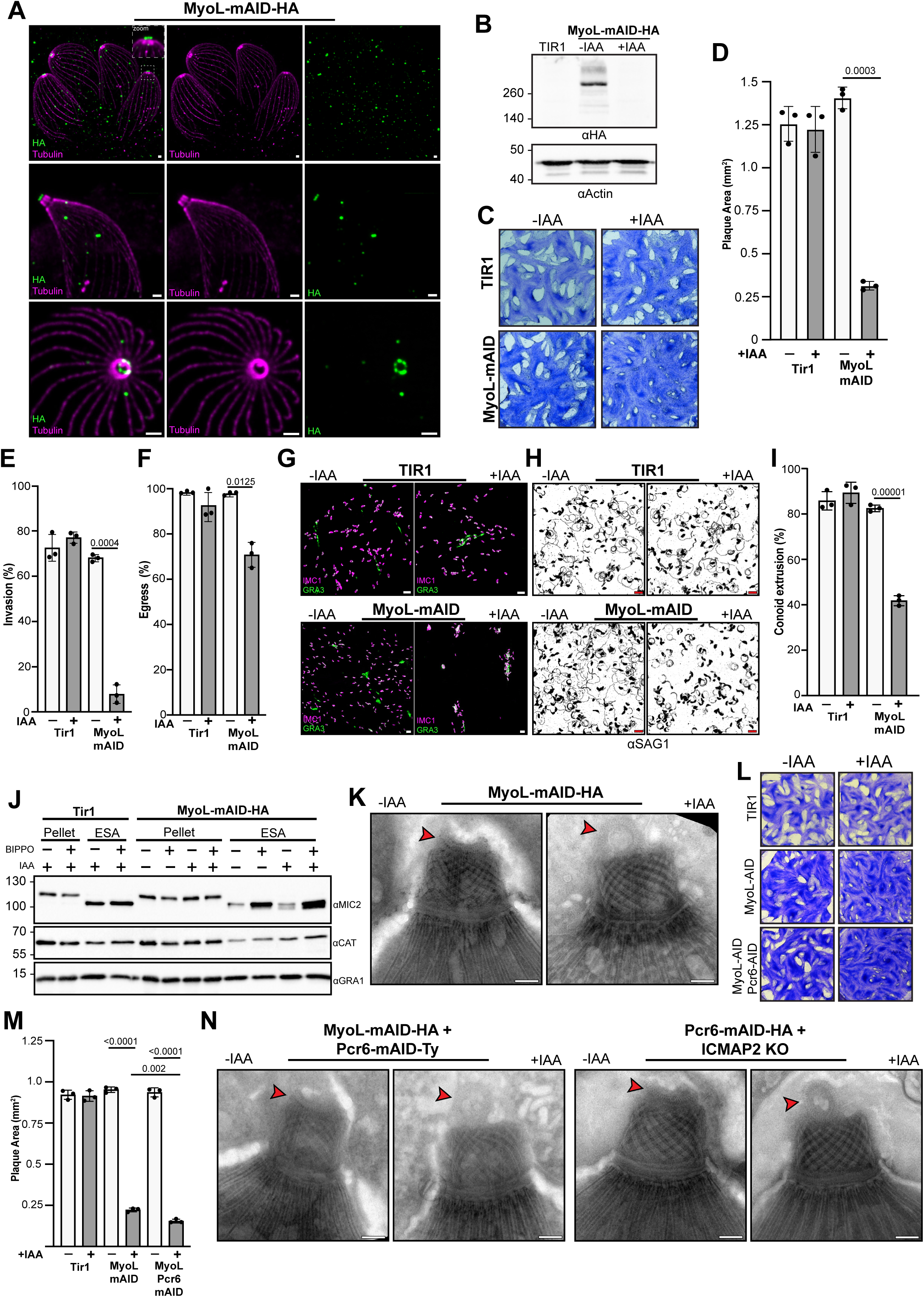
The Myosin-like protein, MyoL, tethers the PCRs to the conoid. **(A)** Localization of the MyoL-mAID-HA protein in U-ExM, both in intracellular and extracellular parasites. Scale bar = 1µm. **(B)** Western blot of MyoL-mAID-HA downregulation upon 24h of IAA treatment, actin is used as the loading control. **(C-D)** Plaque assay images and quantification of plaque area in MyoL depleted parasites. Biological replicates N=3. **(E)** Invasion assay quantification of MyoL depleted parasites. Biological replicates N=3. **(F-G)** Induced-egress assay quantification and images of MyoL depleted parasites. **(H)** Induced-gliding trail assay of MyoL depleted parasites. Scale bar = 25µm. **(I)** Conoid extrusion assay assessed by U-ExM. Biological replicates N=3 **(J)** Induced microneme secretion assay of MyoL depleted parasites assessed by western blot. ESA stands for excretory secretory antigens. **(K)** Negative staining coupled to electron-microscopy of apical complex of MyoL depleted parasites. Scale bar = 200nm. **(L-M)** Plaque assay images and area quantification in MyoL and Pcr6 dually depleted parasites. Biological replicates N=3. **(N)** Negative staining coupled to electron-microscopy of apical complex of MyoL and Pcr6 dually depleted parasites as well as ICMAP2 knock-out and Pcr6 depleted parasites. Scale bar = 200nm.

To elucidate the basis of the gliding defect in MyoL-depleted parasites, negative staining combined with transmission electron microscopy was conducted. In contrast to untreated parasites, which displayed intact apical cytoskeletal structures, MyoL-depleted parasites showed substantial detachment of the PCRs from the conoid **(Fig 5K)**. This defect plausibly explains the impairment gliding motility, as the PCRs integrity is essential for the generation of the apico-basal flux of F-actin needed for parasite motility (Dos Santos Pacheco et al., 2022). Interestingly, Pcr6-depleted parasites displayed a similar phenotype of PCRs detaching from the conoid when analyzed by EM. In addition, the detaching PCRs were observed with a residual attachment at one point to the structure (Dos Santos Pacheco et al., 2022). We hypothesized that MyoL and Pcr6 might function redundantly, with the residual attachment of the PCRs to the conoid being maintained by the non-depleted protein (either Pcr6 or MyoL). To test this, we endogenously tagged Pcr6 with a mAID-Ty cassette in the MyoL-mAID-HA background, generating a double-mAID strain that allows for the simultaneous conditional depletion of both proteins. Correct integration as well as the clonality of the strain was assessed by PCR on genomic and downregulation of each protein was assessed by western blot **(Fig S3 E-F)**. Depletion of MyoL and Pcr6 for seven days lead to severe reduction of plaque smaller than MyoL depleted alone suggesting a possible additive effect **(Fig 5L-M)**. However negative staining on the parasite depleted for these two proteins showed the same phenotype as the single knock down strains, with PCRs detaching from the conoid **(Fig 5N)**. As the ICMTs are tethered to the conoid wall in an asymmetric manner, we wondered if the same attachment point could be used for the residual attachment observed by EM. To assess whether the residual attachment of the PCRs is mediated by the ICMTs, Pcr6 was endogenously tagged with a mAID-HA cassette in an ICMAP2 knockout strain, which results in loss of ICMTs (Dos Santos Pacheco et al., 2024). Successful tagging was verified by PCR on genomic DNA **(Fig S3G)**. Even in absence of the IMCTs, the PCRs kept their point of attachment to the conoid **(Fig 5N)**.

Collectively, these results highlight the critical role of MyoL in the parasite lytic cycle, maintaining the structural integrity of the conoid–PCR connection and thereby enabling efficient motility and dissemination.

### The neck and tail domains of MyoL are sufficient for correct localization and function

To map the functional domains of MyoL and their role in protein localization and function, we designed four truncation constructs and expressed them as a second copy in the UPRT locus of the MyoL-mAID-HA strain **(Fig 6A)**. Proper expression of each construct, as well as the tight downregulation of the endogenous gene were confirmed by western blot **(Fig 6B-C)**. While a wild-type (WT) copy rescued the loss of endogenous MyoL, the N-Ter or Head constructs alone failed to do so **(Fig 6D-E)**. Remarkably, the Neck-Tail construct was able to complement the absence of MyoL to the same level observed for the WT construct **(Fig 6D-E)**. Overexpression driven by the strong tubulin promoter resulted in prominent cytosolic accumulation of all constructs, as observed by IFA **(Fig 6F**). Surprisingly, in presence of the endogenous copy of the MyoL, none of the second-copy constructs could be seen at the apical tip. However, following auxin-induced depletion of the endogenous protein, both the WT and the Neck-Tail constructs were observed accumulating apically **(Fig 6F)**. The WT and Neck-Tail constructs rescued the gliding defect observed upon MyoL depletion **(Fig 6G)** and restored to correct positioning of the PCRs by EM negative staining **(Fig 6H)**. Taken together these results demonstrate that MyoL relies on its neck and tail to properly localize at the PCRs and tether their attachment to the conoid.

**Figure 6.**
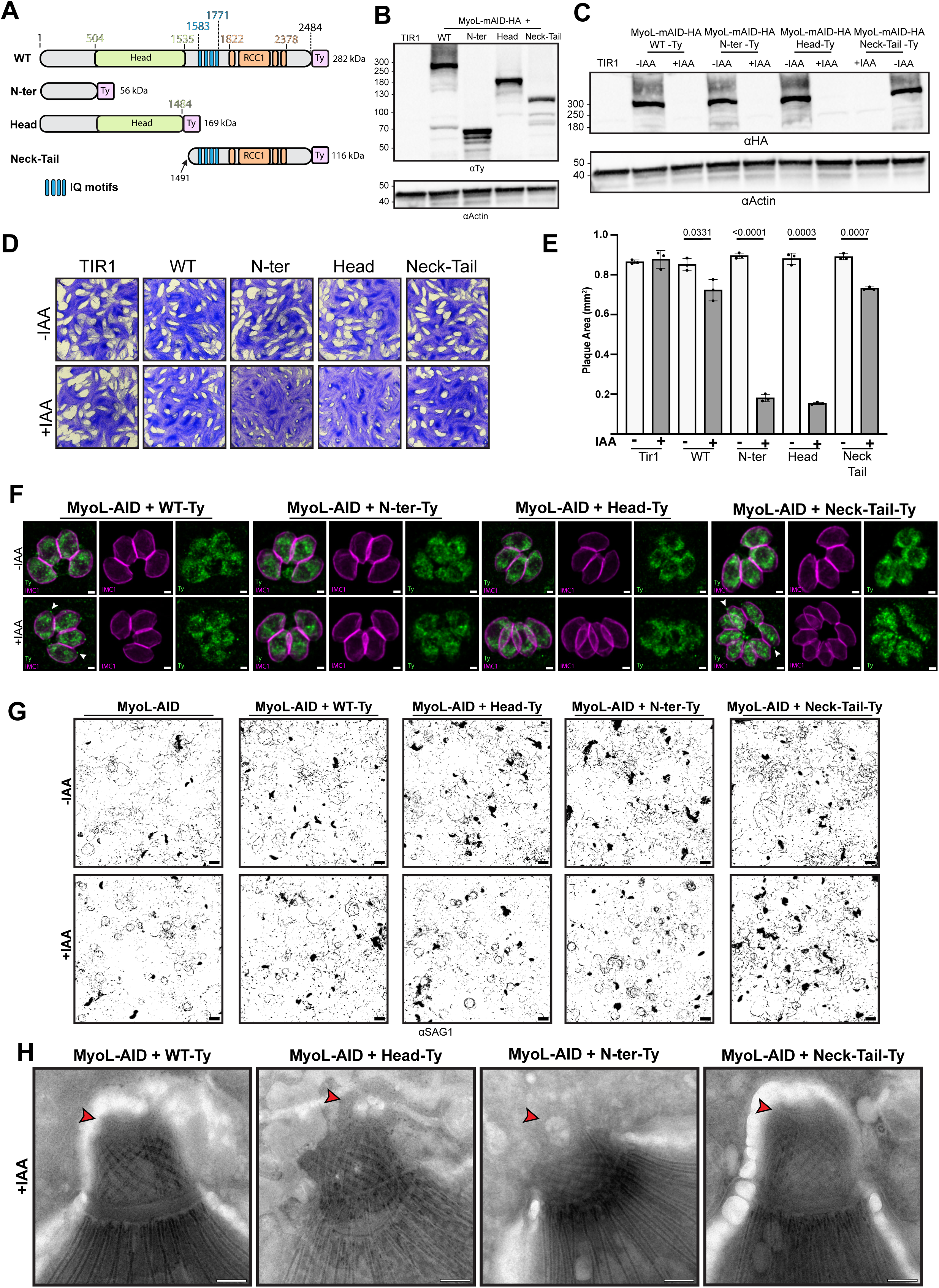
The Neck-Tail of MyoL is responsible for its function and localization. **(A)** Schematic representation of the different MyoL constructs expressed in the UPRT locus of *T. gondii* tachyzoites. **(B)** Western blot showing proper expression of the four constructs expressed in the UPRT locus. **(C)** Western blot showing proper downregulation of MyoL upon addition of auxin in the strains containing the second copy of the MyoL constructs. **(D-E)** Plaque assay images and area quantification in MyoL-mAID-HA strains expressing a second copy of MyoL. Biological replicates N=3. **(F)** IFA images of the different MyoL constructs in intracellular mature parasites. Scale bar = 1µm. **(G)** Induced gliding trail assay of MyoL depleted parasites expressing a second copy of MyoL constructs in the UPRT locus. Scale bar = 25µm. **(H)** Negative staining coupled to electron-microscopy of apical complex of MyoL depleted parasites expressing a second copy of MyoL construct in the UPRT locus. Scale bar = 200nm.

## Discussion

Apicomplexan parasites are defined by their polarized cellular architecture with their apical complex made of cytoskeletal elements and secretory organelles. The conoid plays a pivotal role in regulating the actomyosin based motility and offers a scaffold for the discharge of the secretory organelles critical for invasion. The PCRs represent a conserved conoid-associated structure, with multiple *T. gondii* PCR proteins having orthologs in other apicomplexan species (R Haase et al., 2022). At the structural level, cryo-ET analyses of the PCRs in *Plasmodium falciparum* and *Cryptosporidium parvum* revealed a striking resemblance to those in *T. gondii*, characterized by closely packed repeating units and overlapping densities (Martinez et al., 2023; Sun et al., 2024). To date, two proteins have been identified to be essential for the stability of this conserved structure, Pcr4 and Pcr5 (Dos Santos Pacheco et al., 2022). Both proteins are conserved across the Apicomplexa and more broadly in other alveolates. While their essential role in keeping PCRs integrity was evident in extracellular *T. gondii* tachyzoites (Dos Santos Pacheco et al., 2022), their function was not investigated during daughter cell formation. The PCRs form at an early stage of endodyogeny, preceding most other structures (Padilla et al., 2024). By using other known protein markers of the PCRs, we could monitor the formation of this structure and follow the stability during daughter cell formation. Importantly, our findings suggest that the PCRs can still assemble in the absence of Pcr4/Pcr5, but either cannot fully assemble or undergo disassembly at late stages of daughter cell maturation **(Fig 7A)**. This indicates that Pcr4 and Pcr5 are not required for the initiation of PCRs biogenesis, but rather for their stability and maintenance, implying the existence of other key factors driving their assembly. Amongst these components, there could be daughter cell-restricted proteins, such as Pcr7 (Dos Santos Pacheco et al., 2022) and other proteins found in the structure of the pcr2 ring, like Pcr10 or Pcr12 (pdb: 9AY0) (Jianwei Zeng, 2025). However, depletion of Pcr7 does not impact the formation of the PCRs, suggesting that it is either not contributing to PCRs biogenesis, or it that its function is redundant with another protein. In absence of an observable phenotype upon single-gene downregulation, it remains challenging to identify factors impacting PCRs biogenesis as it would require the implementation of double or multiple gene knock-down strategies. Proximity labeling studies using a daughter-cell restricted protein fusion and coupled with simultaneous depletion of putative candidates involved in PCRs formation could shed light into the biogenesis of this instrumental cytoskeletal structure for the apicomplexan parasites. While revisiting additional PCR-associated proteins, we identified two components, Pcr2 and Pcr3, that remained apically localized even when the overall PCRs structure was disrupted in Pcr4 or Pcr5 depleted parasites. Detailed localization studies, in conjunction with the previously characterized Pcr1, revealed that these proteins adopt a distinctive “half-ring” arrangement, suggesting a non-uniform organization within the PCRs **(Fig 7B**). Moreover, Pcr1, Pcr2 and Pcr3 exhibit differential stability with respect to the PCRs structure, raising questions about their precise subcellular localization. For decades, only two preconoidal rings were described, but the advent of cryo-ET in *T. gondii* revealed the existence of a third preconoidal ring, prompting a re-evaluation of the spatial organization and compartmentalization of these proteins (Gui et al., 2023). Positioned above the two previously characterized PCRs, this third ring is smaller in diameter (Gui et al., 2023). Pcr1, Pcr2 and Pcr3 may belong to this additional ring, potentially accounting for their partial retention following Pcr5 depletion and in case of conoid detachment in RNG2 depleted parasites, although experimental validation of this hypothesis remains to be demonstrated (Haase et al., 2025)

**Figure 7.**
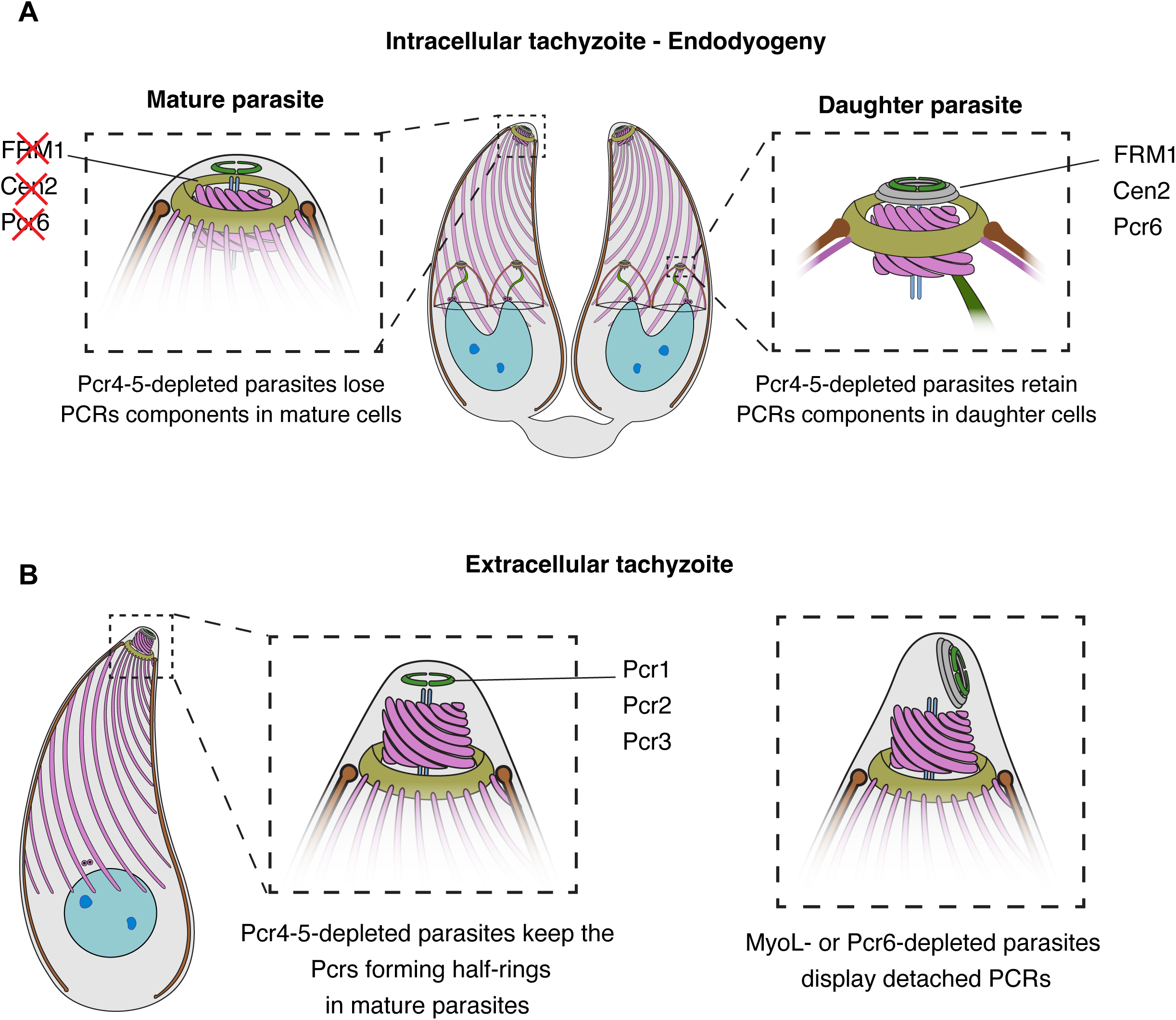
Scheme of the PCRs in several mutants. **(A)** Differential stability of the PCRs and their components between the mature intracellular parasites and the daughter cells parasites in PCR4-5 depleted parasites. **(B)** Left panel shows the maintenance of the half-ring Pcrs, namely Pcr1, Pcr2, and Pcr3, upon Pcr4-5 depletion, suggesting they might be forming the third PCR. Right panel shows the detachment of the PCRs upon MyoL or Pcr6 depletion.

U-ExM led to the unambiguous identification of MyoL as a novel component of the PCRs. Despite lacking the ATP-binding and actin-binding motifs required for motor activity, MyoL serves as a key structural element in tethering the PCRs to the conoid. Functionally, MyoL depletion strongly impaired parasite gliding and host cell invasion, with PCRs detaching in its absence, similar to the effect of Pcr6 downregulation **(Fig 7B)**. This indicates that MyoL acts primarily as a mechanical tether, linking the PCRs to the conoid, rather than as an active myosin motor. Interestingly, both MyoL and Pcr6 display a circular gliding pattern (Dos Santos Pacheco et al., 2022), which was also observed in Rhomboid 4 (ROM4), Myosin A (MyoA), and Myosin Light Chain 1 (MLC1) depleted parasites. ROM4 is a transmembrane protease shedding surface adhesins such as MIC2 to allow proper motility and invasion of the parasite (Buguliskis et al., 2010; Rugarabamu et al., 2015). On the other hand, MyoA and MLC1 are components of the glideosome involved in F-actin translocation towards the basal pole of *T. gondii* tachyzoites (Egarter et al., 2014; Frénal et al., 2010; Herm Götz et al., 2002). Despite their distinct functions, depletion of these three proteins leads to apical accumulation of adhesins, either due to defective processing or impaired translocation to the basal pole. This apical accumulation of adhesins induces a strong attachment of the parasite’s apical tip resulting in circular gliding. In MyoL and Pcr6 mutants, the mispositioning of the PCRs could impair proper translocation of the F-actin flux into the pellicular space, also causing apical accumulation of adhesins and circular gliding. Similarly, *Plasmodium* sporozoites asymmetric secretion of adhesins causes their characteristic circular gliding (Lettermann et al., 2025). Parasites depleted of both MyoL and Pcr6 exhibited smaller plaques than MyoL depletion alone, suggesting an additive effect and possible functional redundancy between the two proteins. However, whether these proteins act together within a complex remains to be determined.

To investigate the functional domains of MyoL, we expressed a second copy of four truncated constructs under the tubulin promoter into the UPRT locus. From the four constructs, only the ones containing the wild-type and the Neck-tail region were able to rescue the phenotype upon MyoL depletion, although not fully.

The constructs localize primarily to the cytosol and are observed at the PCRs only when the endogenous MyoL is depleted, suggesting a saturable binding site for MyoL at this structure. The fragments inserted in the UPRT locus are expressed under the constitutive tubulin promoter and arguably, the change in the promoter could impact in the timing of transcription affecting its expression at a crucial time during early division of the parasite. Also, the endogenous proteins could display some characteristics that are distinct from the complementation fragments that give it a higher affinity towards the PCRs. However, the reason of this partial complementation remains to be elucidated.

The 500 amino acids long N-terminal region preceding the motor domain, which is absent in other myosins, has no predicted structure and no function could be assigned. The motor domain seems to adopt a global conformation reminiscent of myosin heads, but its highly divergent ATP binding site and actin-binding domain suggest that it forms an inactive enzyme. This is further supported by complementation studies showing that a MyoL construct lacking the N-terminal region and the head domain can rescue the phenotype induced by MyoL downregulation. The neck region of MyoL contains 4 predicted IQ motifs that have the potential to mediate binding to MLCs and/ or calmodulins (CaMs). The conoid is populated by three MLCs, namely MLC3/5/7, that associate with Myosin H (MyoH) (Graindorge et al., 2016), as well as by three CaMs, namely CaM1-3, that may interact with MyoH in a calcium-related activation (Long et al., 2017). Similarly, the IQ motifs of MyoL could be involved in the interaction with MLCs or CaMs. In the Cryo-EM structure of the PCR-P2 ring, the N-terminal neck region of MyoL is associated with MLC4, CaM1 and CaM2, identifying at least three myosin light chains associating with MyoL (pdb: 9AY0)(Jianwei Zeng, 2025). The tail domain of MyoL possesses a divergent RCC1 domain predicted to form a seven-bladed propeller structure like RCC1-like domains (RLDs), which have been associated with different functions, including protein-protein and protein-lipid interactions, guanine nucleotide exchange factor and enzyme inhibition (Hadjebi et al., 2008). In contrast to the RCC1 domains of human RCC1 or MyoH, MyoL RLD contains structural motifs intercalated between the blades of the propellers, which may account specific interactions with PCRs or conoid. Arrangement of MyoL head domain in the pcr2 structure suggests that its C-terminal RCC1 domain, which is not defined in the complex structure, may mediate interaction with tubulin at the tip of the conoid. In MyoH, the RCC1-like domain located at the tail region was found to be essential for binding to the conoid tubulin fibers (Graindorge et al., 2016), and it could play a similar role in MyoL. Concordingly, the only fragments that complemented MyoL depletion and correctly localized to the PCRs were the wild-type and the neck-tail constructs, both containing the IQ motifs and the RCC1-like domain. These results indicate that these two elements are essential for MyoL’s function and localization, consistent with other myosins.

Remarkably, both Pcr6 and MyoL are conserved in *Plasmodium* spp. and *Cryptosporidium* spp., suggesting a conserved role as molecular tethers between the conoid and the PCRs in these organisms. Overall, our study advances the understanding of the composition and stability of the PCRs as well as conoid complex assembly in *T. gondii*, a structural element maintained across Apicomplexa and essential for the parasite’s pathogenicity.

## Materials and methods

### Parasite maintenance

*T. gondii* tachyzoites were amplified from HFFs (ATCC) in Dulbecco’s modified Eagle’s medium (DMEM, Gibco) supplemented with 5% fetal calf serum (FCS, Gibco), 2 mM glutamine and 25 µg/ml gentamicin (Gibco). Parasites and HFFs were maintained at 37°C with 5% CO_2_.

### Generation of transgenic strains

All the strains used in this study were constructed using CRISPR-mediated homology directed repair. For construction of 3′-insertional epitope tagging, 40 µg pU6-Universal (pU6-Universal was a gift from Sebastian Lourido, Addgene plasmid # 52694) bearing the gRNA to target the 3′UTR of gene of interest (GOI) was transfected together with PCR product containing homology regions for the selected gene (mAID-HA-HXGPRT, mAID-Ty-DHFR or Ty-DHFR). For enrichment of transfected population, parasites carrying an HXGPRT cassette were selected with 25 mg/ml of mycophenolic acid (MPA) and 50 mg/ml xanthine. Parasites carrying a DHFR cassette were selected with 1 µg/ml of pyrimethamine. In all the assays involving AID-based conditional knockdown system, protein depletion was achieved by addition of 500µM of auxin (IAA) (Brown et al., 2018)

### Genomic PCR to confirm correct integration of phenotyped strains

Genomic DNA of the parasites was extracted using Wizard SV genomic DNA purification system kit. PCR primers were designed to bind outside the 30 bp homology arms in 5′ region as well as to the mAID cassette. TIR1 parasites were used as an independent strain as a PCR control **(Figure S1)**.

### Second copy expression in the UPRT locus for complementation

A codon-remodeled copy of MyoL was synthesized into a modified pUPRT-pTub-G13-Ty plasmid changing the Ty-tag by a 3Ty-tag and (Genscript Company). Specific restrictions sites have been placed within the MyoL sequence allowing excision of the selected regions to generate the according mutants. This synthesized plasmid was name pUPRT-pTub-MyoL-WT-3Ty. To generate the pUPRT-pTub-MyoL-Motor-3Ty the pUPRT-pTub-MyoL-WT-3Ty was digested by EcoRI/XmaI/AvrII (New England Biolabs) allowing the excision of the MyoL gene from the plasmid backbone while in parallel the pUPRT-pTub-MyoL-WT-3Ty was digested by EcoRI/AgeI/HindIII to excise the MyoL fragment containing the motor. They were then ligated using T4 DNA ligase (Thermo Fisher Scientific). To generate the pUPRT-pTub-MyoL-Nter-3Ty the pUPRT-pTub-MyoL-WT-3Ty was digested by EcoRI/XmaI/AvrII allowing the excision of the MyoL gene from the plasmid backbone. In parallel pUPRT-pTub-MyoL-WT-3Ty was digested by EcoRI/XmaI/NdeI (New England Biolabs) to obtain a 7.5kb fragment. This 7.5kb fragment was further digested by BsaWI (New England Biolabs) to obtain the Nter region and ligated with the backbone previously digested as described above. To generate the pUPRT-pTub-MyoL-Tail-3Ty the pUPRT-pTub-MyoL-WT-3Ty was digested by EcoRI/XmaI allowing the excision of the MyoL gene from the plasmid backbone. In parallel the pUPRT-pTub-MyoL-WT-3Ty was digested by MfeI/XmaI (New England Biolabs) allowing the excision of the tail that was further ligated with the plasmid backbone.

For transfection 80μg of plasmid was linearized by NdeI and electroprorated in parasites alongside a plasmid encoding for a 2gRNA targeting the UPRT locus. Selection of positively transfected parasite was performed by adding 5 µM of 5-fluorodeoxyuridine (FUDR) into the media.

### Indirect Immunofluorescence (IFA)

Parasites were fixed using a mix 4% paraformaldehyde, 0,05% Glutaraldehyde during 10 min at room temperature. Following fixation, they were permeabilized by 20-min incubation with PBS-Tx100 0,2% at room temperature. Parasites were blocked during 20 min by PBS-BSA (bovine serum albumin) 5% followed by incubation with the primary antibodies during 1 h in PBS-BSA 2%. Coverslips were washed three times during 5 min by PBS-Tx100 0,2% and then incubated with secondary antibodies diluted in PBS-BSA 2% during 1 h at room temperature. Coverslips were washed three times during 5 min by PBS-Tx100 0,2% and then mounted using Fluoromount G on microscope slides.

### Ultrastructure Expansion Microscopy (U-ExM)

The expansion microscopy protocol applied to *Toxoplasma gondii* tachyzoites was followed as previously described (Dos Santos Pacheco & Soldati-Favre, 2021). Poly-lysine coated coverslips containing extracellular parasites were incubated in PBS containing 0.7% formaldehyde and 1% acrylamide for 3 h at 37°C. Polymerization of expansion gel were performed on ice containing monomer solution (19% sodium acrylate/10% acrylamide/0.1% (1,2-Dihydroxyethylene) bisacrylamide), 0.5% ammonium persulfate (APS) and 0.5% tetramethyl ethylenediamine (TEMED) as described in this study. Fully polymerized gels were denaturated at 95°C for 90 min in the denaturation buffer (200 mM SDS, 200 mM NaCl, 50 mM Tris, pH = 9) and expanded in pure H_2_O overnight. On the next day, the expansion ratio of fully expanded gels was determined by measuring the diameter of gels. Well expanded gels were shrunk in PBS and stained with primary and secondary antibodies diluted in freshly prepared PBS/BSA 2% at 37°C for 2 h. Three washes with PBS/0.1% Tween for 10 min were performed after primary and secondary antibody staining. Stained gels were expanded again in pure H^2^O overnight for further imaging. All U-ExM images used in this study were acquired using Leica TCS SP8 microscopy with the lens HC PL Apo 100x/1.40 Oil CS2. Images were taken with Z-stack and deconvolved with the built-in setting of Leica LAS X or with Huygens software. Final images were processed with ImageJ, and the maximum projected images were presented in this study.

### Electron Microscopy Negative Staining

Electron microscopy of extracellular *T. gondii* tachyzoites was performed as described in (Pacheco et al., 2021). Extracellular parasites were pelleted in PBS. Conoid extrusion was induced by incubation with 40 µl of BIPPO in PBS for 5 min at 37 °C. A 4 µl sample was applied to a glow-discharged 200-mesh Cu electron microscopy grid for 10 min. The excess sample was removed by blotting with filter paper and immediately washing three times in double-distilled water. Finally, the sample was negatively stained with a 0.5% aqueous solution of phosphotungstic acid for 20 s and air-dried. Electron micrographs of parasite apical poles were collected with a Tecnai 20 TEM (FEI, Netherland) operated at 80 kV acceleration voltage equipped with a side-mounted CCD camera (MegaView III, Olympus Imaging Systems) controlled by iTEM software (Olympus Imaging Systems).

### Plaque assay

HFF monolayers were infected with a serial dilution of *T. gondii* tachyzoites and grown for 7 days at 37°C. Cells were fixed with using paraformaldehyde-glutaraldehyde for 10 min followed by a neutralization by PBS/Glycine 0.1M. The fixed monolayer was then stained with crystal violet for 2 h and then washed three times with PBS.

### Invasion assay

Coverslips covered with HFF monolayer were infected with *T. gondii* tachyzoites, centrifuged at 1,000 r.p.m. for 1 min and incubated at 37 °C for 30 min. Cells were then fixed with PFA-GA. Fixed cells were then incubated in 5% PBS-BSA for 20 min of blocking, 1 h with anti-SAG1 antibody and washed three times in PBS. Cells and antibodies were then fixed with 1% formaldehyde for 7 min, permeabilized with 0.2% Triton X100 in PBS for 20 min. Finally, cells were incubated with anti-GAP45 antibodies, washed three times and incubated with appropriate secondary antibodies. Two-hundred parasites were counted for each condition and over three independent biological replicates (*n* = 3).

### Egress assay

Coverslips covered with HFF monolayer were infected with *T. gondii* tachyzoites, centrifuged at 1,000 r.p.m. for 1 min and incubated at 37 °C for 30 h. Cells were then incubated with DMEM media containing BIPPO for 10 min at 37 °C. Cells were then fixed with PFA-GA labelled as described previously, and stained with anti-GAP45 and anti-GRA3 antibodies. Two-hundred vacuoles were counted for each condition in three independent biological replicates (*n* = 3).

### Microneme secretion assay

Freshly egressed parasites were washed twice in warm DMEM media. Parasite pellets were then resuspended in media containing BIPPO and incubated at 37 °C for 10 min. Pellets and supernatant (ESA) were then separated by centrifugation at 2,000*g*. The ESA fraction was then centrifuged at 5,000*g* to remove any cell debris. Pellet fractions were washed once in PBS to remove any ESA remaining. Pellets and ESA were then analyzed by western blot using anti-MIC2, anti-Catalase and anti-GRA1 antibodies. Experiments were performed three times independently (*n* = 3) and a representative replicate is presented in the manuscript. Membrane pictures were acquired using the ImageLab software.

### Gliding trail assay

Freshly egressed tachyzoites were resuspended in warm DMEM media containing BIPPO. Parasites were then put on poly-lysine-coated coverslips, centrifuged at 1,000 r.p.m. for 1 min, and incubated for 10 min at 37 °C. Parasites were then fixed and stained, as described previously in (Dos Santos Pacheco et al., 2022), with anti-SAG1 antibodies (and in absence on Triton X100).

### Conoid extrusion assay

Conoid extrusion was assessed by U-ExM as described before (Dos Santos Pacheco et al., 2022). Briefly, freshly egressed parasites were harvested and conoid extrusion induced by incubating them 15min with warm PBS containing BIPPO. Then, U-ExM was performed and the parasites stained with anti-tubulin antibodies. 200 parasites were counted for each condition, in 3 independent biological replicates.

### Protein depletion assay

HFF monolayers seeded in 6-cm petri dishes were infected with freshly egressed parasites and grown for a 24h in absence or presence of IAA (500 µM). Cells containing the parasites were scrapped and pelleted at 1200 rpm and resuspend in protein loading buffer containing 2% SDS and boiled for 15 min at 95°C. Protein depletion was assessed by Western-Blot using anti-HA antibody against the protein of interest and anti-Actin as a loading control.

## Acknowledgments

We thank the team at the Bioimaging Core Facility, François Prodon, Olivier Brun and Nicolas Liaudet, for their technical assistance and imaging analysis.

## Author contributions

D.S.-F., R.H., A.T.P., N.D.S.P, conceptualized the project. D.S.-F., R.H., A.T.P., O.V., N.D.S.P., conceptualized the methodology and R.H., A.T.P., N.D.S.P, O.V. and B.M., performed the investigations. R.H., A.T.P. performed the formal analysis and R.H wrote the original draft. D.S.-F., R.H., A.T.P., O.V., N.D.S.P, reviewed the manuscript and D.S.F acquired funding and obtained resources. All the authors contributed to this article and approved the submitted version.

## Supplementary Materials

**Supplementary Figure 1.**
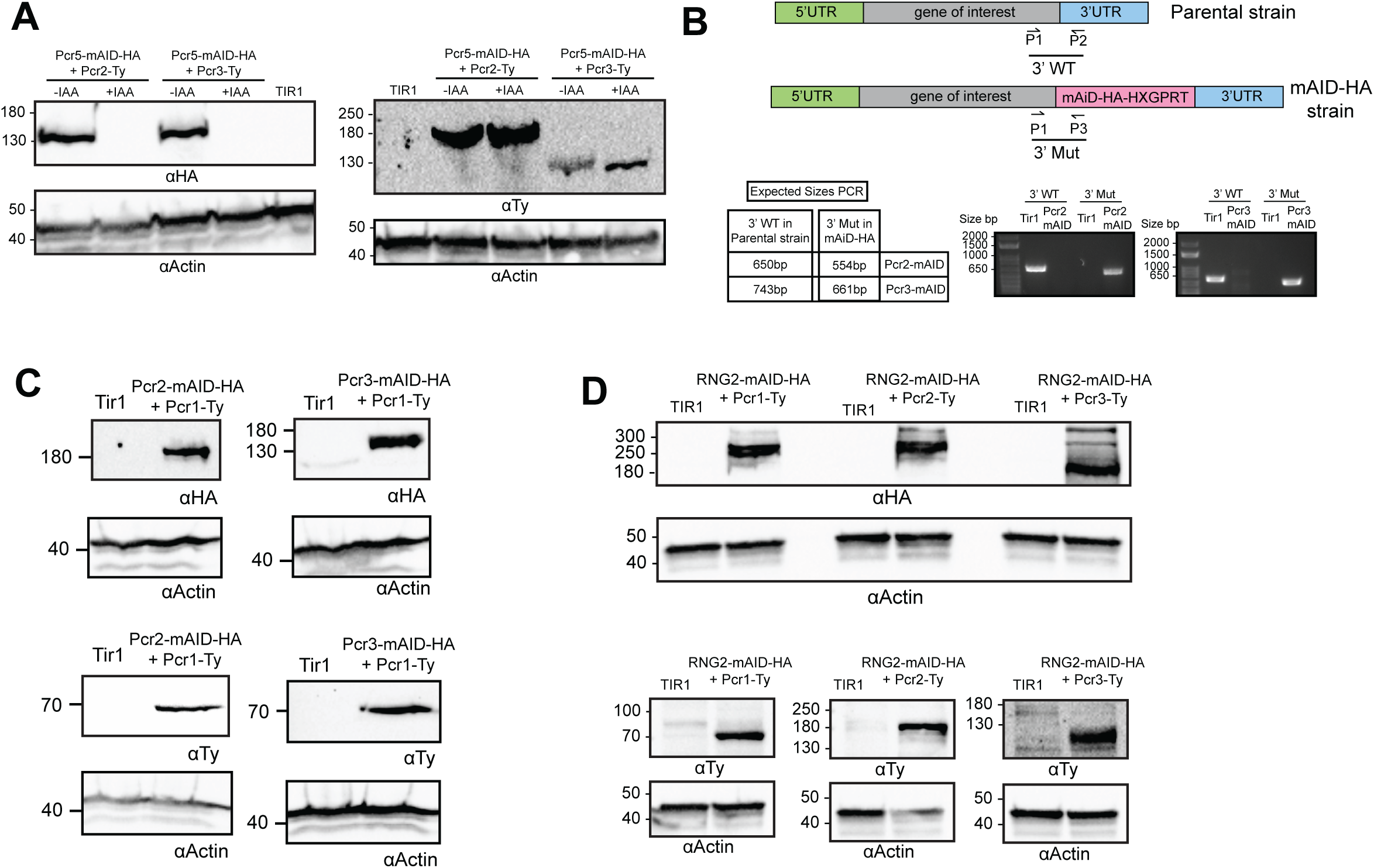
Additional information. **(A)** Western blot analysis of the correct epitope-tagging of Pcr2 and Pcr3 in the Pcr5-mAID-HA background, as well as the proper Pcr5 downregulation upon auxin treatment. **(B)** Integration PCR on genomic DNA to verify the correct integration and clonality of the Pcr2-mAID-HA and Pcr3-mAID-HA strains. **(C)** Western blot analysis of the correct epitope-tagging of the Pcr1 protein in the background of the Pcr2-mAID-HA and Pcr3-mAID-HA strains. **(C)** Western blot analysis of the correct epitope-tagging of the Pcr1, Pcr2 and Pcr3 proteins in the background of the RNG2-mAID-HA strain. **(D)** Western blot analysis of the correct epitope-tagging of the Pcr1, Pcr2 and Pcr3 proteins in the background of the RNG2-mAID-HA strain.

**Supplementary Figure 2.**
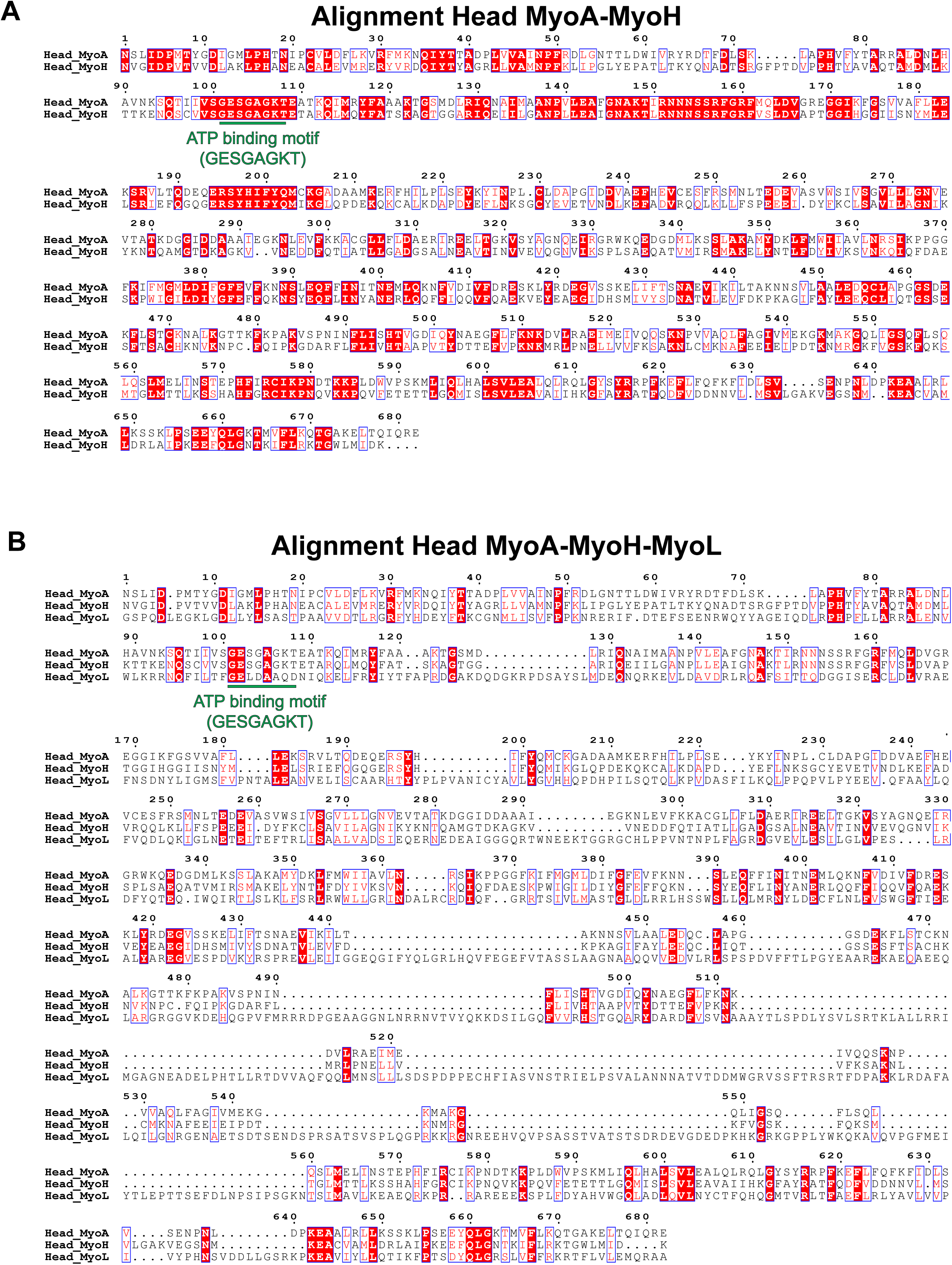
Alignment of the head domain of MyoA, MyoH, and MyoL. **(A)** Sequence alignment of the head domain of MyoA and MyoH, with the ATP binding motif highlighted. **(B)** Sequence alignment of the head domain of MyoA, MyoH, and MyoL, showing poor alignment of MyoL with the two other myosins. The ATP binding motif is also highlighted.

**Supplementary Figure 3.**
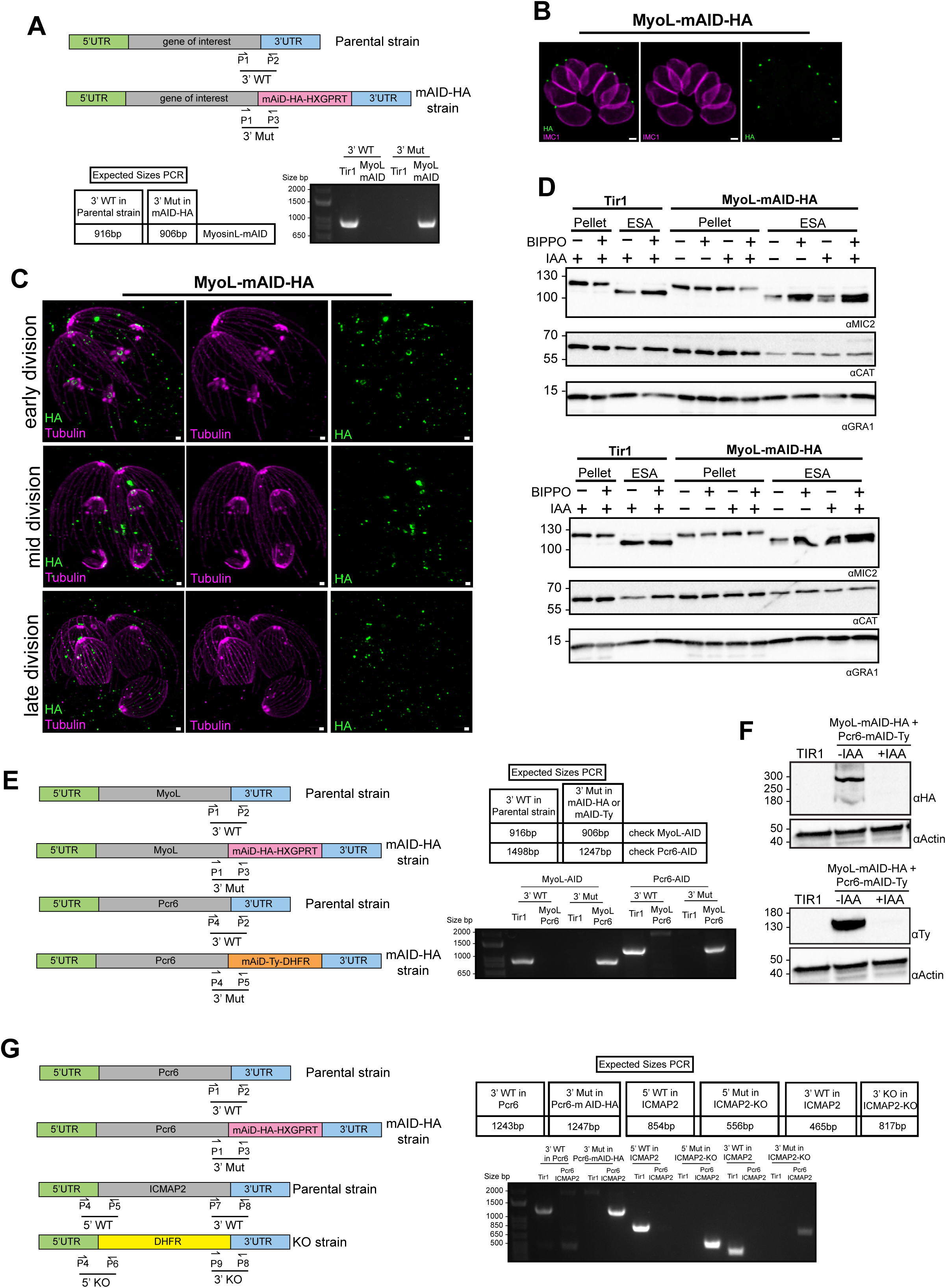
Additional information regarding MyoL phenotyping. **(A)** Integration PCR on genomic DNA to verify the correct integration and clonality of the MyoL-mAID-HA strain. **(B)** IFA showing the apical localization of MyoL in intracellular parasites. Scale bar = 1µm. **(C)** U-ExM images showing different stages of endodyogeny, with MyoL present form the early stages. Scale bar = 3µm. **(D)** Western blot showing the two other biological replicates of microneme secretion. ESA stands for excretory secretory antigens **(E)** Integration PCR on genomic DNA to verify the correct integration and clonality of the double MyoL-mAID-HA + Pcr6-mAID-Ty strain. **(F)** Western blot analysis of the correct depletion of MyoL and Pcr6 in the MyoL-mAID-HA + Pcr6-mAID-Ty strain protein upon 24h of auxin addition. **(G)** Integration PCR on genomic DNA to verify the correct integration and clonality of the double Pcr6-mAID-HA + ICMAP2-KO strain.

**Supplementary Movie 1 – 4. (1-2)** Live microscopy of induced egress of MyoL-mAID-HA parasites, without MyoL depletion. **(3-4)** Live microscopy of induced egress of MyoL depleted parasites.

## References

Adl, S. M., Leander, B. S., Simpson, A. G. B., Archibald, J. M., Anderson, O. R., Bass, D., Bowser, S. S., Brugerolle, G., Farmer, M. A., Karpov, S., Kolisko, M., Lane, C. E., Lodge, D. J., Mann, D. G., Meisterfeld, R., Mendoza, L., Moestrup, Ø., Mozley-Standridge, S. E., Smirnov, A. V., & Spiegel, F. (2007). Diversity, Nomenclature, and Taxonomy of Protists. Systematic Biology, 56(4), 684–689. 10.1080/10635150701494127

Arias Padilla, L. F., Munera Lopez, J., Shibata, A., Murray, J. M., & Hu, K. (2024). The initiation and early development of apical–basal polarity in Toxoplasma gondii. Journal of Cell Science, 137(19). 10.1242/jcs.263436

Brown, K. M., Long, S., & Sibley, L. D. (2018). Conditional Knockdown of Proteins Using Auxin-inducible Degron (AID) Fusions in Toxoplasma gondii. Bio-protocol, 8(4), e2728. 10.21769/BioProtoc.2728

Buguliskis, J. S., Brossier, F., Shuman, J., & Sibley, L. D. (2010). Rhomboid 4 (ROM4) Affects the Processing of Surface Adhesins and Facilitates Host Cell Invasion by Toxoplasma gondii. PLOS Pathogens, 6(4), e1000858. 10.1371/journal.ppat.1000858

Daher, W., Plattner, F., Carlier, M.-F., & Soldati-Favre, D. (2010). Concerted Action of Two Formins in Gliding Motility and Host Cell Invasion by Toxoplasma gondii. PLOS Pathogens, 6(10), e1001132. 10.1371/journal.ppat.1001132

Del Carmen, M. G., Mondragón, M., González, S., & Mondragón, R. (2009). Induction and regulation of conoid extrusion in Toxoplasma gondii. Cellular Microbiology, 11(6), 967–982. 10.1111/j.1462-5822.2009.01304.x

Dos Santos Pacheco, N., Brusini, L., Haase, R., Tosetti, N., Maco, B., Brochet, M., Vadas, O., & Soldati-Favre, D. (2022). Conoid extrusion regulates glideosome assembly to control motility and invasion in Apicomplexa. Nature Microbiology, 7(11), 1777–1790. 10.1038/s41564-022-01212-x

Dos Santos Pacheco, N., & Soldati-Favre, D. (2021). Coupling Auxin-Inducible Degron System with Ultrastructure Expansion Microscopy to Accelerate the Discovery of Gene Function in *Toxoplasma gondii*. In L. M. de Pablos & J. Sotillo (Eds.), Parasite Genomics: Methods and Protocols (pp. 121–137). Springer US.

Dos Santos Pacheco, N., Tell i Puig, A., Guérin, A., Martinez, M., Maco, B., Tosetti, N., Delgado-Betancourt, E., Lunghi, M., Striepen, B., Chang, Y.-W., & Soldati-Favre, D. (2024). Sustained rhoptry docking and discharge requires Toxoplasma gondii intraconoidal microtubule-associated proteins. Nature Communications, 15(1), 379. 10.1038/s41467-023-44631-y

Dos Santos Pacheco, N., Tosetti, N., Koreny, L., Waller, R. F., & Soldati-Favre, D. (2020). Evolution, Composition, Assembly, and Function of the Conoid in Apicomplexa. Trends in Parasitology, 36(8), 688–704. 10.1016/j.pt.2020.05.001

Egarter, S., Andenmatten, N., Jackson, A. J., Whitelaw, J. A., Pall, G., Black, J. A., Ferguson, D. J. P., Tardieux, I., Mogilner, A., & Meissner, M. (2014). The Toxoplasma Acto-MyoA Motor Complex Is Important but Not Essential for Gliding Motility and Host Cell Invasion. PLOS ONE, 9(3), e91819. 10.1371/journal.pone.0091819

Engelberg, K., Bauwens, C., Ferguson, D. J. P., & Gubbels, M.-J. (2025). Co-dependent formation of the *Toxoplasma gondii* subpellicular microtubules and inner membrane skeleton. mBio, 16(9), e01389–01325. 10.1128/mbio.01389-25

Foth, B. J., Goedecke, M. C., & Soldati, D. (2006). New insights into myosin evolution and classification. Proceedings of the National Academy of Sciences, 103(10), 3681–3686. 10.1073/pnas.0506307103

Frénal, K., Jacot, D., Hammoudi, P.-M., Graindorge, A., Maco, B., & Soldati-Favre, D. (2017). Myosin-dependent cell-cell communication controls synchronicity of division in acute and chronic stages of Toxoplasma gondii. Nature Communications, 8(1), 15710. 10.1038/ncomms15710

Frénal, K., Polonais, V., Marq, J.-B., Stratmann, R., Limenitakis, J., & Soldati-Favre, D. (2010). Functional Dissection of the Apicomplexan Glideosome Molecular Architecture. Cell Host & Microbe, 8(4), 343–357. 10.1016/j.chom.2010.09.002

Graindorge, A., Frénal, K., Jacot, D., Salamun, J., Marq, J. B., & Soldati-Favre, D. (2016). The Conoid Associated Motor MyoH Is Indispensable for *Toxoplasma gondii* Entry and Exit from Host Cells. PLOS Pathogens, 12(1), e1005388. 10.1371/journal.ppat.1005388

Guérin, A., & Striepen, B. (2020). The Biology of the Intestinal Intracellular Parasite Cryptosporidium. Cell Host & Microbe, 28(4), 509–515. 10.1016/j.chom.2020.09.007

Gui, L., O’Shaughnessy, W. J., Cai, K., Reetz, E., Reese, M. L., & Nicastro, D. (2023). Cryo-tomography reveals rigid-body motion and organization of apicomplexan invasion machinery. Nature Communications, 14(1), 1775. 10.1038/s41467-023-37327-w

Haase, R., Dos Santos Pacheco, N., & Soldati-Favre, D. (2022). Nanoscale imaging of the conoid and functional dissection of its dynamics in Apicomplexa. Curr Opin Microbiol, 70, 102226. 10.1016/j.mib.2022.102226

Haase, R., Dos Santos Pacheco, N., & Soldati-Favre, D. (2022). Nanoscale imaging of the conoid and functional dissection of its dynamics in Apicomplexa. Current Opinion in Microbiology, 70, 102226. 10.1016/j.mib.2022.102226

Haase, R., Puthenpurackal, A., Maco, B., Guérin, A., & Soldati-Favre, D. (2024). γ-tubulin complex controls the nucleation of tubulin-based structures in Apicomplexa. Molecular Biology of the Cell, 35(9), ar121. 10.1091/mbc.E24-03-0100

Haase, R., Ren, B., Tell i Puig, A., Bonavoglia, A., Marq, J.-B., Visentin, R., Dos Santos Pacheco, N., Maco, B., Mondragón-Flores, R., Vadas, O., & Soldati-Favre, D. (2025). RNG2 tethers the conoid to the apical polar ring in Toxoplasma gondii to enable parasite motility and invasion. PLOS Biology, 23(11), e3003506. 10.1371/journal.pbio.3003506

Hadjebi, O., Casas-Terradellas, E., Garcia-Gonzalo, F. R., & Rosa, J. L. (2008). The RCC1 superfamily: From genes, to function, to disease. Biochimica et Biophysica Acta (BBA) - Molecular Cell Research, 1783(8), 1467–1479. 10.1016/j.bbamcr.2008.03.015

Heintzelman, M. B., & Schwartzman, J. D. (1997). A novel class of unconventional myosins from Toxoplasma gondii. Journal of Molecular Biology, 271(1), 139–146. 10.1006/jmbi.1997.1167

Herm Götz, A., Weiss, S., Stratmann, R., Fujita Becker, S., Ruff, C., Meyhöfer, E., Soldati, T., Manstein, D. J., Geeves, M. A., & Soldati, D. (2002). *Toxoplasma gondii* myosin A and its light chain: a fast, single&#x2010;headed, plus&#x2010;end&#x2010;directed motor. The EMBO Journal, 21(9), 2149–2158. 10.1093/emboj/21.9.2149

Hu, K., Roos, D. S., & Murray, J. M. (2002). A novel polymer of tubulin forms the conoid of Toxoplasma gondii. Journal of Cell Biology, 156(6), 1039–1050. 10.1083/jcb.200112086

Jacot, D., Tosetti, N., Pires, I., Stock, J., Graindorge, A., Hung, Y.-F., Han, H., Tewari, R., Kursula, I., & Soldati-Favre, D. (2016). An Apicomplexan Actin-Binding Protein Serves as a Connector and Lipid Sensor to Coordinate Motility and Invasion. Cell Host & Microbe, 20(6), 731–743. 10.1016/j.chom.2016.10.020

Jianwei Zeng, Y. F., Pengge Qian, Wei Huang, Qingwei Niu, Wandy L. Beatty, Alan Brown, L. David SIbley, Rui Zhang. (2025). Atomic models of the Toxoplasma cell invasion machinery. Nature structural and molecular biolofy

Koreny, L., Zeeshan, M., Barylyuk, K., Tromer, E. C., van Hooff, J. J. E., Brady, D., Ke, H., Chelaghma, S., Ferguson, D. J. P., Eme, L., Tewari, R., & Waller, R. F. (2021). Molecular characterization of the conoid complex in *Toxoplasma* reveals its conservation in all apicomplexans, including *Plasmodium* species. PLOS Biology, 19(3), e3001081. 10.1371/journal.pbio.3001081

Lettermann, L., Singer, M., Steinbrück, S., Ziebert, F., Kanatani, S., Sinnis, P., Frischknecht, F., & Schwarz, U. S. (2025). Chirality of malaria parasites determines their motion patterns. Nature Physics. 10.1038/s41567-025-03096-0

Long, S., Brown, K. M., Drewry, L. L., Anthony, B., Phan, I. Q. H., & Sibley, L. D. (2017). Calmodulin-like proteins localized to the conoid regulate motility and cell invasion by *Toxoplasma gondii*. PLOS Pathogens, 13(5), e1006379. 10.1371/journal.ppat.1006379

Martinez, M., Mageswaran, S. K., Guérin, A., Chen, W. D., Thompson, C. P., Chavin, S., Soldati-Favre, D., Striepen, B., & Chang, Y.-W. (2023). Origin and arrangement of actin filaments for gliding motility in apicomplexan parasites revealed by cryoelectron tomography. Nature Communications, 14(1), 4800. 10.1038/s41467-023-40520-6

Mondragon, R., & Frixione, E. (1996). Ca2+-Dependence of Conoid Extrusion in *Toxoplasma gondii* Tachyzoites. Journal of Eukaryotic Microbiology, 43(2), 120–127. 10.1111/j.1550-7408.1996.tb04491.x

Monteiro, V. G., de Melo, E. J. T., Attias, M., & de Souza, W. (2001). Morphological Changes during Conoid Extrusion in Toxoplasma gondii Tachyzoites Treated with Calcium Ionophore. Journal of Structural Biology, 136(3), 181–189. 10.1006/jsbi.2002.4444

Montoya, J. G., & Liesenfeld, O. (2004). Toxoplasmosis. The Lancet, 363(9425), 1965–1976. 10.1016/S0140-6736(04)16412-X

Munera Lopez, J., Tengganu, I. F., Liu, J., Murray, J. M., Arias Padilla, L. F., Zhang, Y., Brown, P. T., Florens, L., & Hu, K. (2022). An apical protein, Pcr2, is required for persistent movement by the human parasite Toxoplasma gondii. PLOS Pathogens, 18(8), e1010776. 10.1371/journal.ppat.1010776

Pacheco, N. D. S., Tosetti, N., Krishnan, A., Haase, R., Maco, B., Suarez, C., Ren, B., & Soldati-Favre, D. (2021). Revisiting the Role of Toxoplasma gondii ERK7 in the Maintenance and Stability of the Apical Complex. mBio, 12(5), 10.1128/mbio.02057-02021. https://doi.org/doi:10.1128/mbio.02057-21

Padilla, L. F. A., Murray, J. M., & Hu, K. (2024). The initiation and early development of the tubulin-containing cytoskeleton in the human parasite Toxoplasma gondii. Molecular Biology of the Cell, 35(3), ar37. 10.1091/mbc.E23-11-0418

Phillips, M. A., Burrows, J. N., Manyando, C., van Huijsduijnen, R. H., Van Voorhis, W. C., & Wells, T. N. C. (2017). Malaria. Nature Reviews Disease Primers, 3(1), 17050. 10.1038/nrdp.2017.50

Pospich, S., Sweeney, H. L., Houdusse, A., & Raunser, S. (2021). High-resolution structures of the actomyosin-V complex in three nucleotide states provide insights into the force generation mechanism. eLife, 10, e73724. 10.7554/eLife.73724

Ren, B., Haase, R., Patray, S., Nguyen, Q., Maco, B., Dos Santos Pacheco, N., Chang, Y.-W., & Soldati-Favre, D. (2024). Architecture of the Toxoplasma gondii apical polar ring and its role in gliding motility and invasion. Proceedings of the National Academy of Sciences, 121(46), e2416602121. 10.1073/pnas.2416602121

Rugarabamu, G., Marq, J.-B., Guérin, A., Lebrun, M., & Soldati-Favre, D. (2015). Distinct contribution of Toxoplasma gondii rhomboid proteases 4 and 5 to micronemal protein protease 1 activity during invasion. Molecular Microbiology, 97(2), 244–262. 10.1111/mmi.13021

Sidik, S. M., Huet, D., Ganesan, S. M., Huynh, M.-H., Wang, T., Nasamu, A. S., Thiru, P., Saeij, J. P. J., Carruthers, V. B., Niles, J. C., & Lourido, S. (2016). A Genome-wide CRISPR Screen in *Toxoplasma* Identifies Essential Apicomplexan Genes. Cell, 166(6), 1423–1435.e1412. 10.1016/j.cell.2016.08.019

Sun, S. Y., Segev-Zarko, L.-a., Pintilie, G. D., Kim, C. Y., Staggers, S. R., Schmid, M. F., Egan, E. S., Chiu, W., & Boothroyd, J. C. (2024). Cryogenic electron tomography reveals novel structures in the apical complex of *Plasmodium falciparum*. mBio, 15(4), e02864–02823. 10.1128/mbio.02864-23

Tosetti, N., Dos Santos Pacheco, N., Soldati-Favre, D., & Jacot, D. (2019). Three F-actin assembly centers regulate organelle inheritance, cell-cell communication and motility in *Toxoplasma gondii*. eLife, 8, e42669. 10.7554/eLife.42669

